# Panakeia - A universal tool for bacterial pangenome analysis

**DOI:** 10.1101/2021.03.02.433540

**Authors:** Sina Beier, Nicholas R. Thomson

## Abstract

We present Panakeia, a tool which strives to be easy to use and providing a detailed view of the pangenome structure which can efficiently be utilised for discovery, or further in-depth analysis, of features of interest. It analyses synteny and multiple structural patterns of the pangenome, giving insights into the biological diversity and evolution of the studied taxon. Panakeia hence provides both broad and detailed information on the structure of a pangenome, for diverse and highly clonal populations of bacteria. Development of new pan-genome analysis tools is important, as the pangenome of a microbial species has become an important method to define the diversity of a selected taxon, most commonly a species, in the last years. This enables comparison of strains from different ecological niches and can be used to define the functional potential in a bacterial population. It gives us a much better view of microbial genomics than can be gained from singular genomes which after all are just single representatives of a much more varied population.

Previously published pangenome tools often reduce the information to a presence/absence matrix of unconnected genes or generate massive hard to interpret output graphs. However, Panakeia includes synteny and structural information and presents it in a way that can readily be used for further analysis. Panakeia can be downloaded at https://github.com/BioSina/Panakeia.

## 1 Introduction

Pangenome analysis is increasingly popular, especially in microbiology, where the concept of species can be blurry at best [1, 2] and isolated single genomes are are of limited value for understanding evolution and population diversity of microbes. With this interest in microbial populations comes the need to understand the complete genetic makeup from the individual isolates to the whole genus and more. The accumulated genome of these groups of samples is called a pangenome [4].

Several new tools have emerged in the last years to provide automated analysis pipelines. Each of them has a different focus into which functions of the pangenome are to be studied. Many investigate the size and members of common for the taxon compared to rarely found genes, generally called the core and accessory pangenome. In the last years, including analysis of syntenic structures in the pangenome has risen in importance, as these structures often include interesting functional operons, or provide important contextual information e.g. evidence of having been horizontally acquired. However, in large pangenomes, there can be many such loci or structures, and this makes it hard to determine which are essential and relevant for further investigation. Often, this information is presented in huge pangenome graph structures. These get increasingly complicated with growing number of input genomes and the inherent diversity in these genomes. These complexities can make it time-consuming and tedious for the user to interpret the results from pangenome analysis.

Here we describe Panakeia, an analysis pipeline for prokaryotic pangenomes which includes a graphic representation of the pangenomewith a special focus on analyzing the synteny and specific genomic patterns found in the dataset. Hence, Panakeia enables a detailed look into the structure of a pangenome, compared to the overview look of core and accessory genes that pangenome analysis tools generally provide. The syntenic structures Panakeia identifies can be filtered and divided into groups which hint at their function and origin. The graph structure also inherently offers multiple ways to detect interesting structural patterns.

Panakeia is designed to be used together with the Pantagruel [12] pipeline for the reconstruction of the evolutionary history of all genes in the dataset. Pantagruel uses phylogenetic algorithms to determine the evolutionary history of genes and gene families. It can differentiate between evolutionary events and horizontal gene transfer and indicate if a gene transfer event has occurred between different lineages of the same species.

Together, the pangenome structure, including the synteny and patterns identifying small variants as well as large structural variants between the genomes determined by Panakeia and the genomic history of the genes determined by Pantagruel gives a detailed view of the evolutionary history and genomic plasticity of a set of closely related prokaryotic genomes.

## 2 Methods

Panakeia is written in Python 3, utilising the NetworkX [3] package for graph generation and analysis. It is split into a preprocessing script for clustering, the main pipeline and multiple post-processing scripts.

### 2.1 Preprocessing

Input genomic DNA sequences for Panakeia should be annotated in GFF3 format, which is a standard output format for many prokaryotic annotation tools like Prokka [6] or PGAP [8]. For the following clustering step, the predicted protein sequences have to be available, either through the output of the annotation or by extracting them from the annotated genome using other tools. If annotation of the genomes is not feasible, Panakeia can be run without functional annotation of the genomes but predicted protein sequences have to be provided. They can be determined using fast protein prediction methods like Prodigal [5].

To analyse the pangenome, the proteins have to be clustered using the provided script *Clustering.py*, which takes a single fastA file with all predicted protein sequences from the genomes and the number of input genomes as input. All proteins from the annotated genomes are clustered using cd-hit [7] in an iterative process, going from 90 % sequence similarity required to cluster proteins over 80% similarity and 75% down to 70 %. Clusters which include at least the number of input genomes are kept, while proteins in smaller clusters are re-clustered with a lower similarity threshold until the threshold of 70 % sequence similarity is reached. At this point, all remaining clusters are kept. One random sequence is chosen as a representative for each cluster.

### 2.2 Pan-genome analysis

Panakeia reads in the genome annotations for each input genome in GFF3 format, generating so-called strain graphs, which are graphs using proteins as nodes and connecting local neighbours with edges. Another type of edge is added for paralogs, which are detected by reading in the clusters generated by the previous step and defining each pair of proteins which are from the same genome and classified into the same cluster as paralogs.

The clusters are also used as nodes for a pangenome graph, which is similar to the strain graphs but has whole clusters as nodes (annotated by the features of the clusters representative sequence). Edges are added to the pangenome graph if two clusters include neighbouring proteins in at least one of the input genomes. Thus the edges determine the synteny information for the pangenome graph. Edge weight is determined by the number of genomes in which the connected nodes are neighbouring each other. Pangenome nodes also include information on the maximum number of paralogs predicted proteins from the cluster can have in one single genome.

These paralogous clusters are then subdivided into subclusters by determining the minimal number of unrelated neighbourhoods the cluster is part of in all genomes. Each neighbourhood generates a subcluster, with the original cluster nodes being removed from the graph. Subcluster nodes share the same representative, but the features determining in which strains they occur and if they belong to the pangenome core or accessory proteome are updated for each subcluster.

### 2.3 Associating proteins with chromosomal information

If finished or nearly finished genomes which are assembled in the correct number of expected contigs (chromosomes or plasmids) are available, the pangenome graph can be annotated with basic structural information using the *ChromosomizeAll.py* and *ChromosomizePangenome.py* scripts. Clusters which have a member found in one of the sets of complete genomes are assigned to a chromosome, undetermined (if they were present on either chromosome/plasmid) and unknown (if the cluster was not present in any of the complete assemblies). Chromosomal information is added to the clusters in the pangenome graph and all proteins (determined by their cluster membership) in the strain graphs using the information from the complete genomes. This process will further be called ’chromosomizing’ the genome.

### 2.4 Determining structural patterns in the pangenome

Panakeia can detect different types of patterns in the pangenome graph using the *Patterns.py* script, which potentially correspond with specific biological features. The detected patterns are

- orphans
- uniques
- variants
- insertions
- indels

The implied meaning of these patterns is further described in Section 3.2.3. The detected patterns are returned as text-based files - either lists or tables, depending on the type of pattern - and as separate graph files which includes only the occurrences of each pattern. This makes it possible to either look at them separately, or overlay the pattern information onto the full pangenome graph by using the *MarkPattern.py* script which uses one of the pattern graphs to highlight the pangenome graph.

### 2.5 Highlighting external information

If phylogenetic information or other information clustering or grouping the input genomes - for example lineages defined by Pantagruel or groups defined through metadata features - are available to the user, groups of genomes can be highlighted onto the pangenome graph by using the *HighlightStrains.py* script. This will highlight proteins and neighborhood relations occurring in a list of strains. Proteins and edges already included in the pangenome graph will be marked in orange. Proteins and edges which did not reach the threshold for minimal weight in the pangenome graph will be added and highlighted in yellow. This enables the user to depict the part of the pangenome and functional potential covered by a defined group of genomes and also extract a group-specific sub-pangenome more easily using network visualisation tools.

### 2.6 Inclusion of Pantagruel output

Panakeia is designed to work closely together with the Pantagruel [12] pipeline for reconstruction of gene histories in bacterial pangenome datasets. Pantagruel output includes information on phylogenetic clades for the input genomes and genes specifically present in these clades or specifically absent compared to neighbouring clades. This presence and absence information informs about evolutionary changes of gene uptake or loss in defined phylogenetic clades which might be related to the fitness in a specific environmental niche or the virulence of pathogenic genomes. The *HighlightClades.py* script uses the Pantagruel output files, specifically the species tree clade definition file and a version of a clade-specific gene set file including either the specifically present or absent genes, to highlight all proteins from the genomes in the clade and mark either the specifically present or absent proteins. This highlighting connects the synteny information to the evolutionary history of these groups of genes.

## 3 Results

### 3.1 Running Pankeia

To prepare for running Panakeia, the user will need the protein sequences extracted from annotation or protein prediction for all the input genomes in a single multi-fastA file, as well as the matching GFF3 files with protein annotations for each separate genome. To produce the protein clusters, the protein fastA will be used as input for clustering the protein sequences with the provided clustering script. This script returns a file containing all cluster information and another file with all the representative sequences for each cluster, including all the functional annotations, if they have been provided in the input GFFs. The main Panakeia pipeline then requires selection of an output directory for all results, an input directory which has to include all GFF3 files from the input genomes, the clustering output file and the file with the representative sequences as they are provided by *Clustering.py*.

Additionally, parameters can be provieded defining the percentages required to identify a protein as belonging to the different partitions which compromise the pangenome: hard core, soft core and shell. Hard core represents proteins which strongly belong to the backbone of the investigated taxon and should not be missing from any genomes. Soft core proteins occur in most input genomes, but are not essential to the taxon, so might be missing occasionally. Shell proteins occur in groups of genomes, and potentially include the structure which differentiate between lineages or functional groups of organisms. Finally, the left over proteins belong to the cloud, which includes only proteins which occur rarely or even in singular genomes. The cloud often includes erroneous protein predictions caused by assembly errors, but also new horizontal gene transfers.

The default values for these parameters are set as 0.99 (meaning a protein has to occur in 99% of the genomes to be counted, acoounting for some proteins missing due to sequencing and assembly errors) for hard core, 0.95 to count it into soft core and 0.15 to count it to the shell. Everything occurring in less than 15 % of the genomes will, in this case, be counted as belonging to the cloud. It is also possible to set a parameter defining the minimal number of occurrences in the pangenome for an edge (local neighbourhood connection between to proteins) to be drawn. This defaults to drawing all edges. For large numbers of input genomes it can be helpful to set this parameter to remove edges which only occur in single genomes or very small numbers of genomes. This automatically reduces the prevalence of assembly errors and generates a more simplified version of the pangenome graph.

### 3.2 Panakeia output

#### 3.2.1 Genome graphs

Panakeia generates a so-called *strain graph* for each input genome, which is provided in the GraphML format that can be loaded into many commonly available graph visualisation tools, including Cytoscape [11]. The strain graphs can then be chromosomised of finished or high-quality reference genomes are available amongst the input genomes to help define the different chromosomes and/or plasmids. An example for a chromosomised version of a *Vibrio cholerae* genome is provided in Figure 1 and 2.

**Figure 1:**
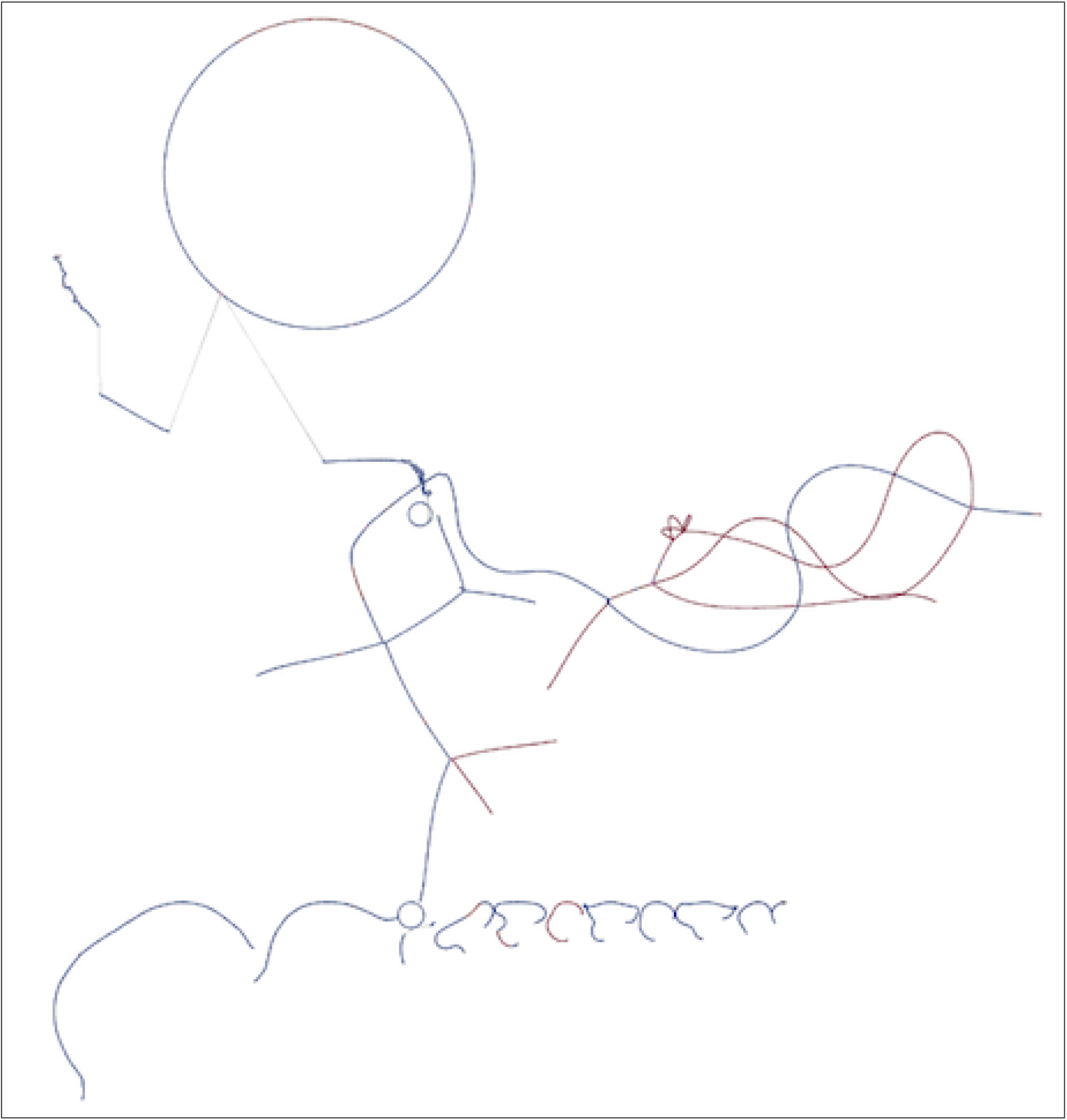
chromosomised view of a strain graph for one *V. cholerae* genome. Vibrio species generally have two chromosomes. Chromosome 1 is marked blue, chromosome 2 is marked red, proteins, where the chromosome cannot be decided as they occur on both chromosomes for the chromosomal reference genomes, are marked pink and proteins which cannot be assigned a chromosome as they do not occur in the references are grey. The proteins are connected to their local neighbours on the contigs, but also their paralogs in the genome. Hence different contigs show a connection.

**Figure 2:**
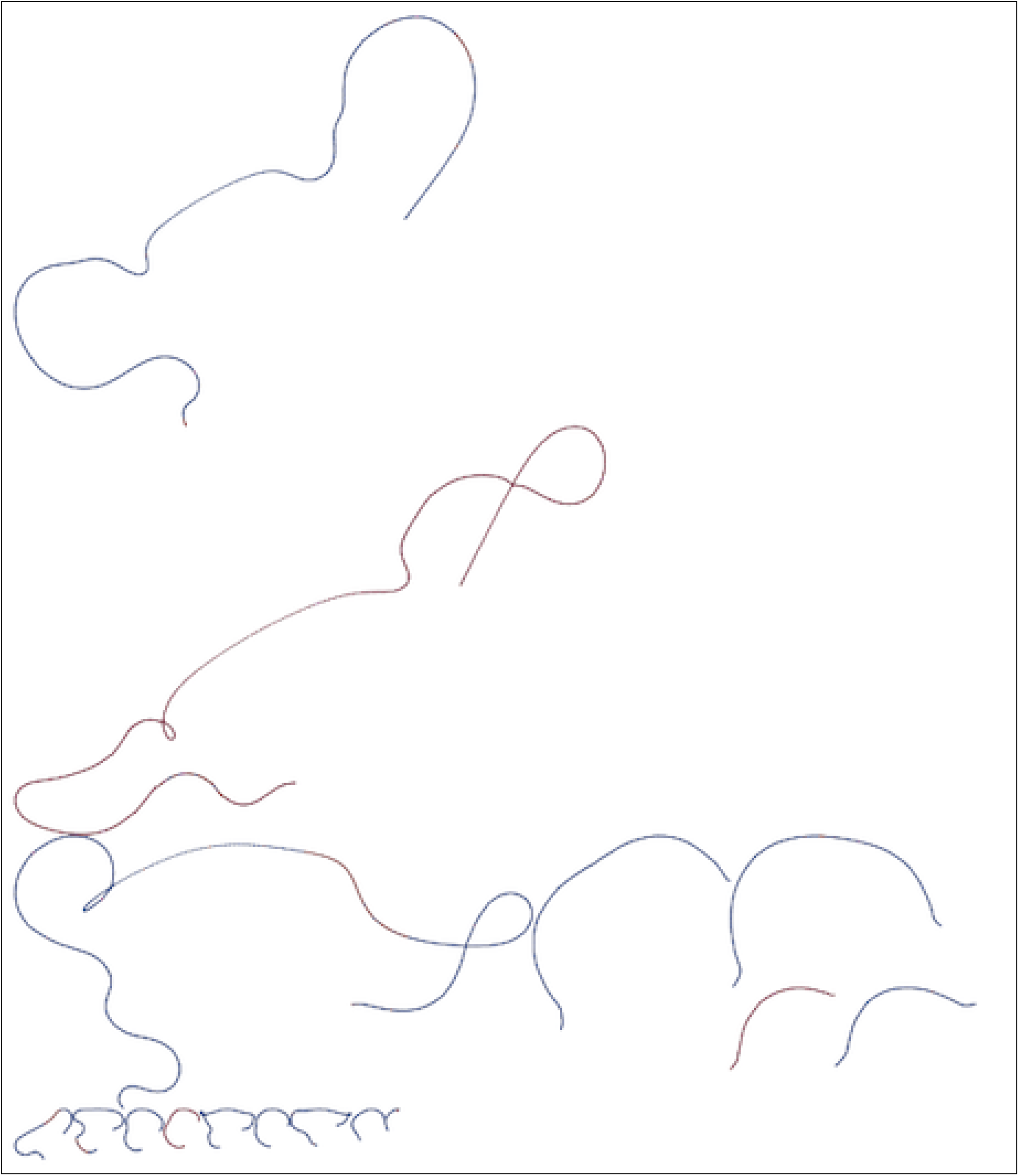
chromosomised view of a strain graph for one *V. cholerae* genome. Vibrio species generally have two chromosomes. Chromosome 1 is marked blue, chromosome 2 is marked red, proteins, where the chromosome cannot be decided as they occur on both chromosomes for the chromosomal reference genomes, are marked pink and proteins which cannot be assigned a chromosome as they do not occur in the references are grey. The proteins are connected to their local neighbours on the contigs, but not by their paralogs. Hence this is a graph of the proteins as they are located on the contigs.

Strain graphs help to assess the assembly of an isolate, as assembly errors caused by repetitive sequences often show up as loops of paralogous proteins in the synteny of the strain graph. Assembly errors caused by single paralogs or small mobile elements are easily found in chromosomised strain graphs as they have a change of chromosome/plasmid assignment around a single or a group of proteins which occur on both chromosomes, as shown in Figure 3.

**Figure 3:**
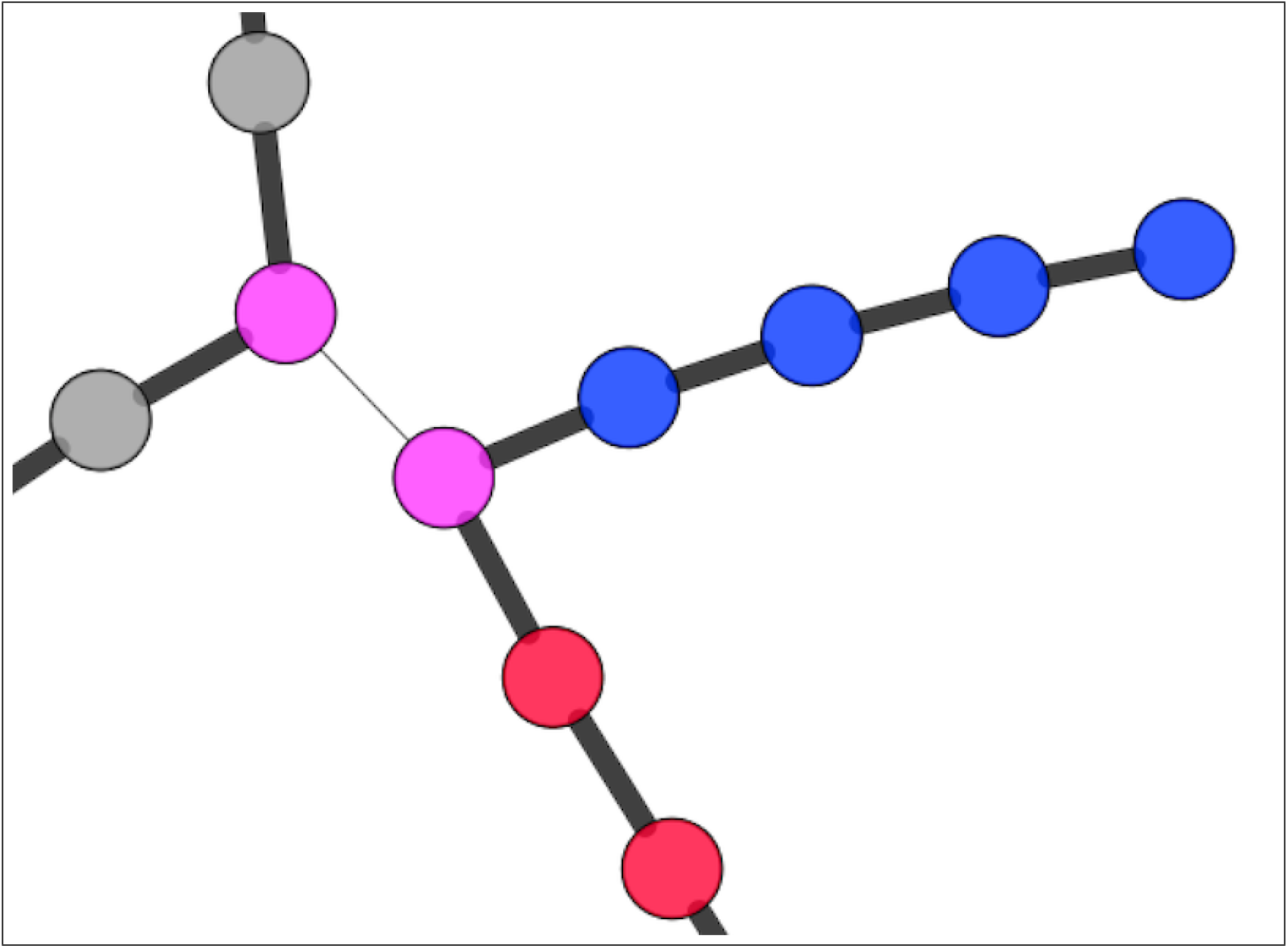
A misassembly in a chromosomised strain graph, as visible by the blue proteins belonging to chromosome 1 and red proteins belonging to chromosome two, connected by a pink (undecided) protein which is annotated as an IS200/IS605 family transposase. The wide black edges mark local neighbourhood, the thin grey edge marks paralogous relation of two proteins.

Strain graphs also serve as input for further processing like *HighlightStrains.py*, which can be used to highlight one or more strains as an overlay over the complete pangenome, and scripts specific to include output of the Pantagruel [12] pipeline into the pangenome graph.

#### 3.2.2 Pangenome graph and analysis

The full pangenome graph is the most important output of the Panakeia pipeline, it enables all further analyses steps and visualisation of the data. This graph includes all protein clusters and all local edges for neighborhood connections that occur in more genomes than the given minimal support threshold. The nodes hold information like the strains they occur in if they are hard core, soft core, accessory or cloud, the representative protein of the cluster, the maximal number of paralogs in a genome, number of proteins in the cluster and a weighting based on the average number of proteins in this cluster per genome. If available, it also includes the functional annotation of the cluster, and if the pangenome has been chromosomised previously, it includes the chromosomal prediction. An example of a chromosomised pangenome graph is shown in Figure 4.

**Figure 4:**
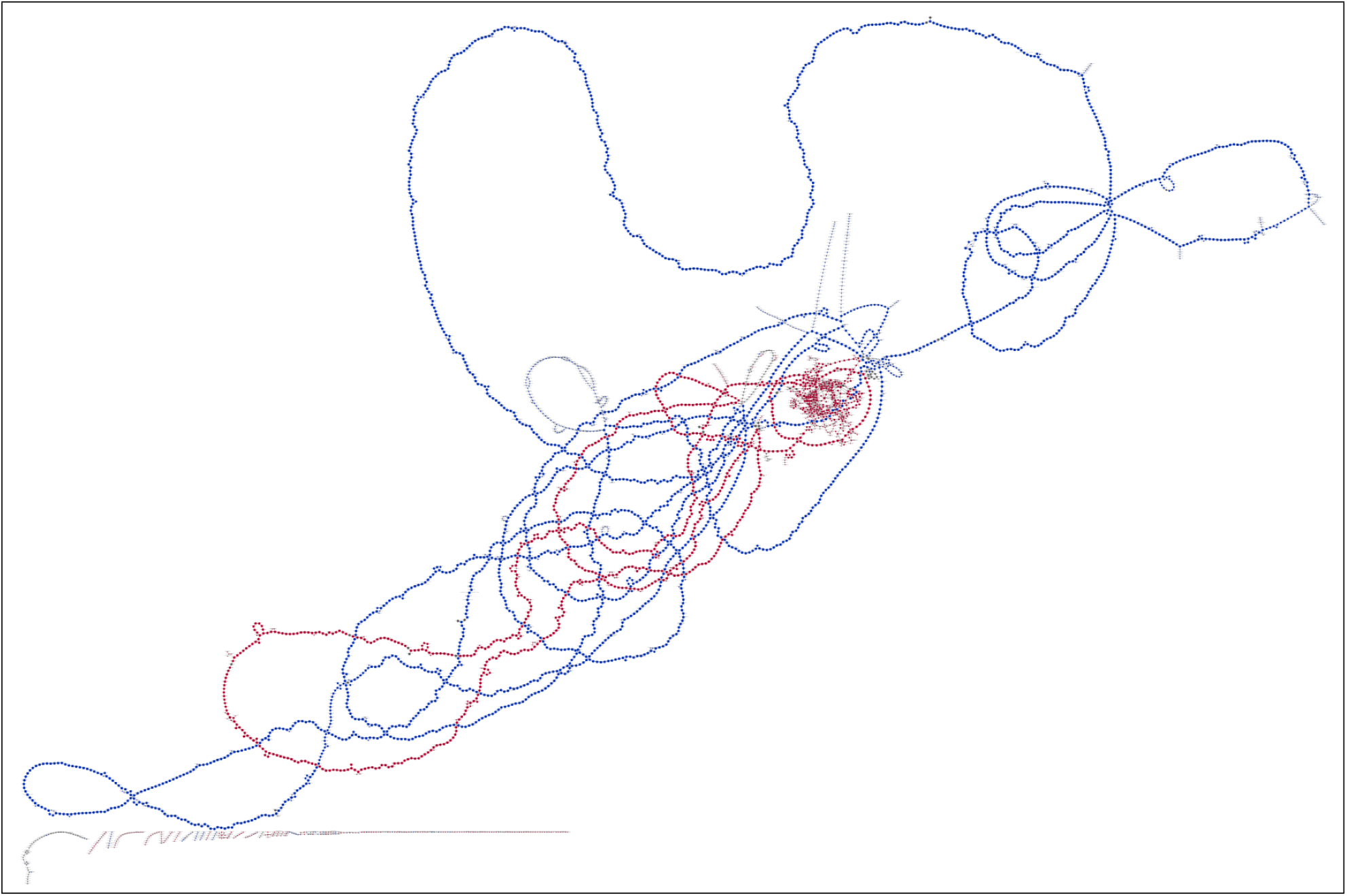
chromosomised view of a pangenome graph for 42 finished *V. cholerae* as found on RefSeq. As the species has two chromosomes, chromosome 1 is marked blue, and chromosome 2 is marked red, proteins, where the chromosome cannot be decided as they occur on both chromosomes for the chromosomal reference genomes, are marked dark grey and proteins which cannot be assigned a chromosome as they do not occur in the references are light grey.

#### 3.2.3 Pattern extraction

After the pangenome graph is generated and annotated, biologically relevant patterns can be detected using the *Pattern.py* script, as described in Section 2.4.

Orphans are proteins which do not have a common placement in the genomes. They are either caused by contigs from contamination of the genomic sample with external DNA, assembly problems around this specific gene or highly variable locations in different genomes. The latter can hint at the proteins belonging to small mobile elements. The functional annotation can help differentiate between these cases.

’Uniques’ are proteins which only occur in a single genome. They are either caused by contamination or annotation errors or are truly novel, perhaps as a result of recent horizontal gene transfer into that genome. This is when combining information from Pantagruel can be especially useful because it can be used to further investigate their evolutionary history.

’Variants’ are proteins which have multiple amino acid variants occuring in the same genomic neighborhood. This only occurs if the sequence variation is bigger than the cutoff used for clustering. They can be variants of the same functional gene with small changes in the amino acid sequence, but also include truncated proteins or pseudogenes which have lost their function in a part of the genomes.

’Insertions’ stand for rare short insertions into one or a few genomes. Much like the Uniquess, they can be caused by contamination but also can hint at horizontal gene transfer or incorporation of a plasmid or other mobile genetic element.

’InDels’ are common insertions or rare deletions (relative to the number of input genomes) that are found in the pangenome. These are potentially structural variants and hint at gene loss in a few of the genomes or at large-scale homologous recombination events, which can be typically seen around capsule or O-antigen gene clusters.

The *Pattern.py* script can also generate a tabular representation of the presence/absence matrix for all proteins and genomes in the dataset. Patterns extracted from the pangenome are saved either in tabular or text form, depending on the pattern. They are also saved as a graph only containing the occurrences of the pattern for visualization. An example of an extracted InDel graph is shown in Figure 5. The matching text output includes a tab-delimited list of clusters for each InDel, displayed in one InDel per line. Text output for variants is similar, with a list of the varying different protein clusters in the same structural location per line.

**Figure 5:**
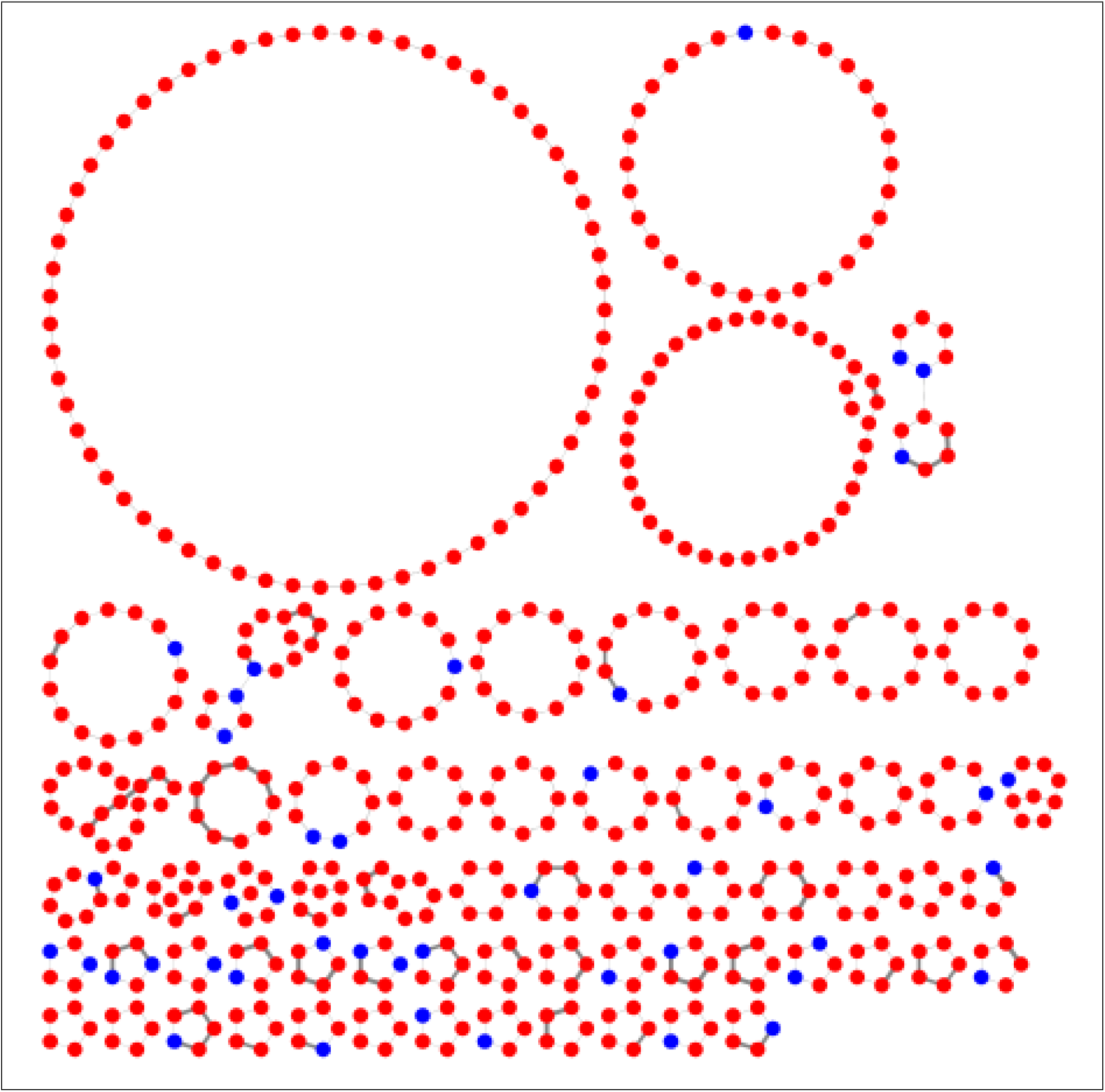
InDels extracted from a set of 318 genomes. The core proteins are marked blue, the non-core proteins in red. InDels are generally non-core (very rare deletions are not detected). They are often “attached” to the core through one or two core proteins which mark the location of the inDel. In this case, the inDel is either inherited in most genomes or a mobile element specific to a single insertion position.

Text output for insertion sequences is in tabular form, including the node degree (number of attached edges) and node weight for each potential insertion sequence.

### 3.3 Comparison to other pangenome tools

Panakeia uses a simple approach to clustering proteins compared to other pangenome tools and focuses on using the synteny and paralogy information from the input genomes to increase the available information content and accuracy of the resulting pangenome graph. Synteny information is also used by over novel tools like Panaroo [10] and PPanGGOLiN [9], which do cluster genes instead of proteins. In general, pangenome tools often specialise by either in showing an overview of the full functional diversity of a studied taxon or by correcting small errors caused by technical or algorithmic restrictions and giving information on a long list of small detailed structures. Both approaches output can be cumbersome and need prior knowledge to interpret and utilise the information appropriately.

Panakeia incorporates no default correction steps other than splitting paralogous clusters and enabling the user to set a cutoff for minimum support of edges, but it can give specific insights by using the various patterns described in Section 3.2.3 together with the functional annotation. This makes it possible to move from a broad view of the available diversity to details about interesting structures and patterns which define the difference between groups of genomes and can lead to helpful insights about the evolution and adaptation of these organisms.

To explain the differences of Panakeia to other pangenome tools, here we will compare it to two relatively current developments in the field and define the different use cases for each of the tools.

PPanGGOLiN is named after the <u>P</u>artitioned <u>P</u>an<u>G</u>enome <u>G</u>raph <u>O</u>f <u>L</u>inked <u>N</u>eighbors it uses to analyse the pangenome. It focuses on partitioning the homologous gene clusters it uses into a persistent (core), cloud and one or more shell partitions. Panakeia in comparison divides the core into hard and soft core to be able to track novel gene loss which has only occurred in a small subset of the taxons phylogeny through using the pattern detection to detect InDels in the backbone of the pangenome. Panakeia does not further divide the shell - which is usually the largest group of protein clusters for large and diverse datasets - because these clusters are the most important to be able to detect the patterns associated with features of a group of genomes. Dividing the shell into further groups would potentially restrict the size of a group of genomes which include one of these patterns for them to be found. Gautreau *et al.* [9] mention the possibility of predicting genomic islands in the shell and cloud as a future option of PPaNGOLiN, Panakeia already includes the functionality through the pattern search.

Panaroo focuses on correcting the inclusion of genes into the pangenome, filtering out potential contamination and genes classified as being based on misassembly or annotation errors. This leads to very clean pangenome graphs even for large input datasets but could result in the loss of new horizontal gene transfer events or changes in the structure caused by the movement of mobile elements. Hence this approach might not be appropriate for taxa with known or predicted high diversity, high genome plasticity or even just for datasets where more fragmented assemblies have to be included. By using the parameter to set a minimum weight for edges to be included in the pangenome graph, Panakeia can be set up to filter out many similar errors easily, as they are generally creating rare or even unique edges. The potential errors will show up as singletons in the pangenome graph and can then be detected by the pattern search. Many visualisation tools will also allow you to detect the singletons as nodes of degree 0 and remove them from the graph for a cleaner visualisation, making it unnecessary to already include this detection when building the pangenome graph.

## 4 Discussion

Panakeia is designed to combine pan-genome analysis with genomic information from isolate genomes to improve the information collected both about the studied population and the genome plasticity and structure of single members in it. Additionally, it can be linked to phylogenetic and evolutionary information provided by Pantagruel. This combination will help to improve our understanding of how microbial genomes function, adapt and evolve. Automatically extracting and highlighting interesting patterns enables researchers to focus on potentially interesting features of the genomes without the need of painstakingly finding them in long lists, tables and huge graphs.

Based on Python 3 with minimal extra packages and standardised input and output file formats, the pipeline is widely applicable and easy to install, maintain and use.

The pipeline is not optimised for speed, but aimed at reducing the manual work necessary to find patterns of interest in large pangenome datasets, which in turn reduces the time needed to extract useful information from these datasets. The information about the patterns can be exported both in text-based files which are easy to use in further analysis and as GraphML files for visualisation and manual curation of the results. Hence, Panakeia is a tool which is usable in many settings and for people from different backgrounds and with different requirements for their work.

## References

[1] L-M Bobay and H Ochman. Biological Species Are Universal across Life’s Domains Genome Biology and Evolution, 2017, volume 9(3), pages 491–501

[2] M Vos. A species concept for bacteria based on adaptive divergence Trends in Microbiology, 2011, volume 19(1), pages 1–7

[3] A Hagberg et al. Exploring Network Structure, Dynamics, and Function using NetworkX Proc. SciPy 2008, pages 11–15

[4] D Medini et al. The microbial pangenome Curr. Opin. Genet. Dev., 2005, volume 15(6), pages 589–594

[5] D Hyatt et al. Prodigal: prokaryotic gene recognition and translation initiation site identification. BMC Bioinformatics, 2010, volume 11(1), page 119

[6] T Seemann. Prokka: rapid prokaryotic genome annotation Bioinformatics, 2014, issue 14, pages 2068–2069.

[7] W Li and A Godzik. Cd-hit: A fast program for clustering and comparing large sets of protein or nucleotide sequences Bioinformatics, 2006, volume 22, issue 13, pages 1658–1659.

[8] T Tatusova et al. NCBI prokaryotic genome annotation pipeline Nucleic Acids Res., 2016 Aug 19, volume 44, issue 14, pages 6614–24.

[9] G Gautreau et al. PPanGGOLiN: Depicting microbial diversity via a Partitioned Pangenome Graph PLoS Comput Biol, 2020 volume 16(3): e1007732

[10] G Tonkin-Hill et al. Producing Polished Prokaryotic Pangenomes with the Panaroo Pipeline BioRxiv, 2020, https://doi.org/10.1101/2020.01.28.922989

[11] ME Smoot et al. Cytoscape 2.8: new features for data integration and network visualization Bioinformatics, 2011, volume 27, pages 431–432.

[12] F Lassalle et al. Automated Reconstruction of All Gene Histories in Large Bacterial Pangenome Datasets and Search for Co-Evolved Gene Modules with Pantagruel visualization bioRxiv, 2019, 586495.

